# Mechanical properties of the stigmatic cell wall mediate pollen tube path in Arabidopsis

**DOI:** 10.1101/384321

**Authors:** Lucie Riglet, Frédérique Rozier, Chie Kodera, Isabelle Fobis-Loisy, Thierry Gaude

## Abstract

Successful fertilization in angiosperms depends on the proper trajectory of pollen tubes through the pistil tissues to reach the ovules. Pollen tubes first grow within the cell wall of the papilla cells, applying pressure to the cell. Mechanical forces are known to play a major role in plant cell shape by controlling the orientation of cortical microtubules (CMTs), which in turn mediate deposition of cellulose microfibrils (CMFs). Here, by combining cell imaging and genetic approaches, we show that isotropic reorientation of CMTs and CMFs in aged and *katanin1-5* (*ktn1-5*) papilla cells is accompanied by a tendency of pollen tubes to coil around the papillae. Furthermore, we uncover that aged and *ktn1-5* papilla cells have a softer cell wall and provide less resistance to pollen tube growth. Our results reveal an unexpected role for KTN1 in pollen tube guidance by ensuring mechanical anisotropy of the papilla cell wall.

## INTRODUCTION

Following deposition of dehydrated pollen grains on the receptive surface of the female organ, the stigma, pollen rehydrates, germinates and produces a pollen tube that carries the male gametes toward the ovules where the double fertilization takes place. This long itinerary through the different tissues of the pistil is finely controlled, avoiding misrouting of the pollen tube and hence assuring proper delivery of the sperm cells to the female gametes. In *Arabidopsis thaliana*, pollen tubes grow within the cell wall of papillae of the stigmatic epidermis, and then through the transmitting tissue of the style and ovary (Lennon and Lord, 2000). The transmitting tissue has an essential function in pollen tube guidance, providing chemical attractants and nutrients (Crawford and Yanofsky, 2008; Higashiyama and Hamamura, 2008). In contrast to these accumulating data showing the existence of factors mediating pollen tube growth in the pistil, whether guidance cues exist at the very early stage of pollen tube emergence and growth in the papilla cell wall remains largely unknown. The cell wall constitutes a stiff substrate and hence a mechanical barrier to pollen tube progression. There are numerous examples in animal cells demonstrating that mechanical properties of the cellular environment, and in particular rigidity, mediate cell signalling, proliferation, differentiation and migration (Discher et al., 2005; Ermis et al., 2018; Fu et al., 2010). In plant cells, cell wall rigidity depends mainly on its major component, cellulose, which is synthesized by plasma membrane-localized cellulose synthase complexes (CSCs) moving along cortical microtubule (CMT) tracks (Paredez et al., 2006). While penetrating the cell wall, the pollen tube exerts a pressure onto the stigmatic cell (Sanati Nezhad and Geitmann, 2013). Such physical forces are known to reorganise the cortical microtubules (CMTs), which by directing CSCs to the plasma membrane, reinforce wall stiffness by novel cellulose microfibril (CMF) synthesis (Paredez et al., 2006; Sampathkumar et al., 2014). Hence, there is an intricate interconnection between CMT organisation, CMF deposition and cell wall rigidity (Xiao et al., 2016; Xiao and Anderson, 2016). A major regulatory element of CMT dynamics is the KATANIN (KTN1) microtubule-severing enzyme, which allows CMT reorientation following mechanical stimulation (Louveaux et al., 2016; Sampathkumar et al., 2014; Uyttewaal et al., 2012). Here we investigated whether the CMT network of papilla cells might contribute to pollen tube growth and guidance in stigmatic cells by combining cell imaging techniques, genetic tools and atomic force microscopy. We show that isotropic reorientation of CMTs occurs in aged Col-0 and *ktn1-5* papilla cells, which is accompanied by a change in the growth direction of pollen tubes that tend to make coils on papillae. We demonstrate that both papilla types exhibit softer cell walls that correlates with a faster growth rate of pollen tubes. Besides, we show that CMT reorganisation is associated with isotropic rearrangement of CMFs. To determine whether the coiled phenotype of pollen tubes could be a general feature of papillae affected in cytoskeleton or cell wall composition, we tested a series of cell wall mutants including the *xxt1 xxt2* double mutant that displays cytoskeleton orientation defects and reduced stiffness of the cell wall. Remarkably, none used as female induces coiled growth of wild-type pollen tubes. Altogether, our results indicate that cytoskeleton dynamics and mechanical properties of the cell wall that both depend on KTN1 activity have a major role in guiding early pollen tube growth in stigma papillae.

## RESULTS

### CMT dynamic pattern and pollen tube growth during stigma development

To assess the functional role of stigmatic CMTs in pollen – papilla cell interaction, we first analysed their organisation in papillae at stages 12 to 15 of stigma development as described (Smyth et al., 1990) (Fig. 1*A* and *B*). We generated a transgenic line expressing the CMT marker MAP65.1-citrine under the control of the stigma specific promoter SLR1 (Fobis-Loisy et al., 2007). Before (stage 12) and at anthesis (stage 13), the CMTs were aligned perpendicularly to the longitudinal axis of papilla cells and were highly anisotropic (median value of 0.40) (Fig. 1*C* and *D*). At stage 14, when anthers extend above the stigma, the CMT pattern became less organised, with a higher variability in anisotropy values. Finally, at stage 15 when the stigma extends above anthers, CMT anisotropy had a median value of 0.09 indicative of an isotropic orientation of CMTs (Fig. 1*C* and *D*). These findings reveal that the papilla CMT cytoskeleton is dynamic during development, with a change of CMT array orientation from anisotropy to isotropy. We then wondered whether this change in CMT organisation could be correlated with pollen tube growth. To this end, we self-pollinated Col-0 papillae from stages 12 to 15 and examined pollen tube growth one hour after pollination by scanning electron microscopy (SEM) (Fig. 2*A*). At stage 12 and 13, we found most (∼60%) pollen tubes to grow straight in the papillae, whereas about 30% and 10% of tubes made half-turn or one turn around stigmatic cells, respectively (Fig. 2*B*). At later stages of development, the tendency to coil around the papillae dramatically increased, with more than 35% of pollen tubes at stage 14 and more than 55% at stage 15 making one or more than one turn around papillae. These results suggest that CMT organisation in the papilla impacts the direction of growth of pollen tubes and that loss of CMT anisotropy is associated with coiled growth.

**Fig. 1.**
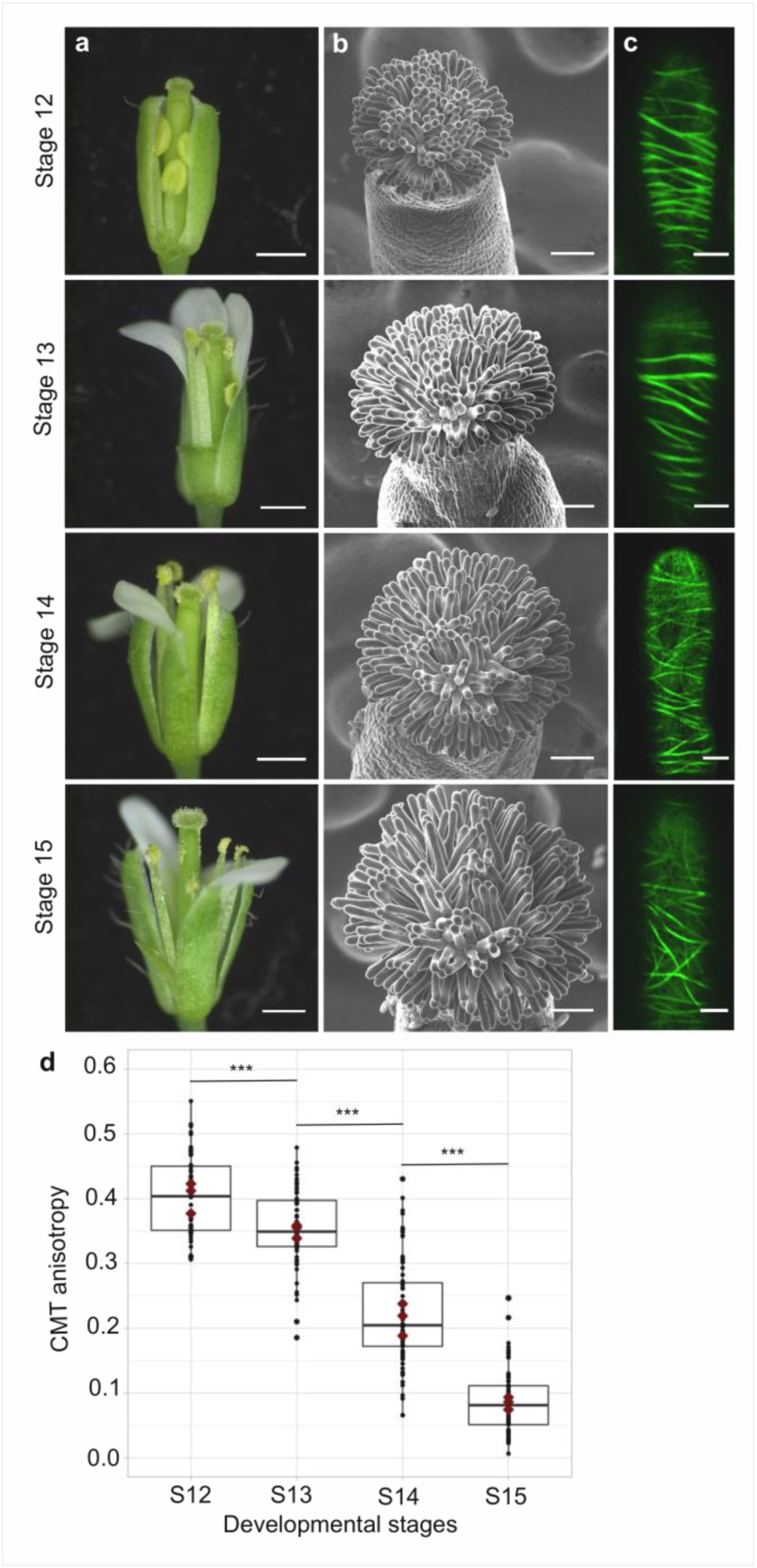
CMT organisation during papilla cell development. (*A*) Flower development of *A. thaliana* from developmental stages 12 to 15. Scale bar, 500 μm. (*B*) Upper view of the stigma during development by SEM. Scale bar, 50 μm. (*C*) Confocal images of papilla cells expressing MAP65-citrine at each stage of development. Scale bar, 5 μm. (*D*) Quantitative analysis of CMT array anisotropy of papilla cells from stages 12 to 15. The red dots correspond to the mean values of the three replicates. Statistical differences were calculated using a Shapiro-Wilk test to evaluate the normality and then a Wilcoxon test, ***P < 0.01. N > 4 stigmas, n > 60 papilla cells for each stage.

**Fig. 2.**
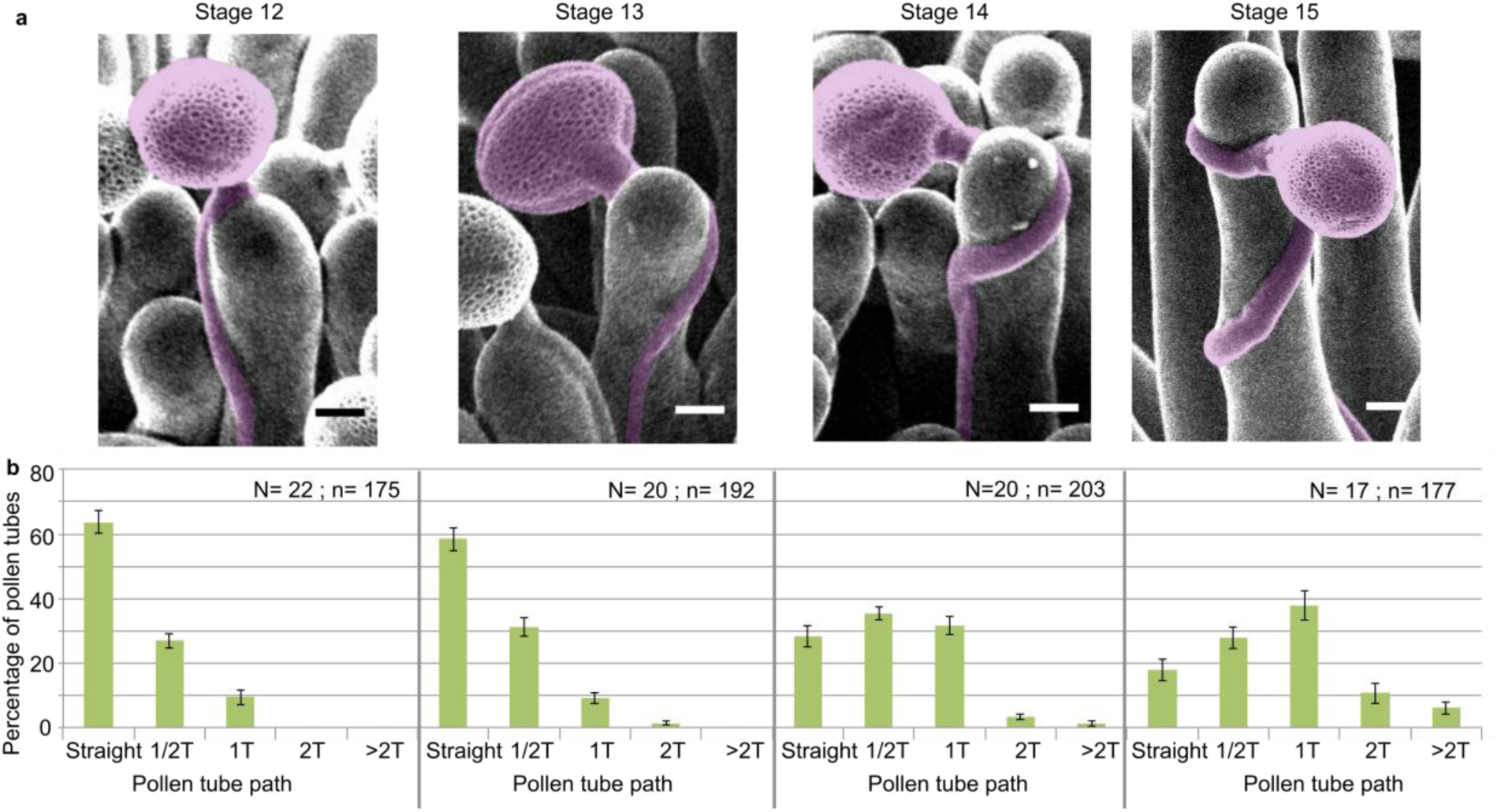
Pollen tube growth behaviour on papillae during development. (*A*) SEM images of Col-0 papillae pollinated with Col-0 pollen, one hour post pollination, from stages 12 to 15; pollen and tubes were artificially colorised. Scale bar, 5 μm. (*B*) Quantification of the number of turns (T) made by the pollen tube on papillae from stages 12 to 15. Data are expressed as mean +/-s.e.m. A Chi-Square test for independence (at 6 degree of freedom) was used to compare all stages and demonstrated that the number of turns was significantly different between stages (***P < 0.01). N corresponds to the number of stigmas analysed in this study and n, the number of papillae.

### Impaired CMT dynamics of papillae affects pollen tube growth direction

To confirm the relation between stigma CMTs and pollen behaviour, we examined pollen tube growth on stigmas of the *katanin1-5* mutant, which is known to exhibit reduced CMT array anisotropy in root cells (Bichet et al., 2001; Burk et al., 2001; Burk and Ye, 2002). Because the CMT organisation in *ktn1-5* papillae is unknown, we crossed *ktn1-5* with the MAP65-1-citrine marker line and found that CMT arrays were more isotropic in *ktn1-5* papillae when compared with those of the WT (Fig. 3*A* and *C*). Using SEM, we then analysed Col-0 pollen behaviour on *ktn1-5* stigmatic cells at stage 13 (Fig. 3*B*). We observed that Col-0 pollen tubes acquired a strong tendency to coil around *ktn1-5* papillae, with above 60% of tubes making one or more than one turn around papillae, sometimes making up to 6 turns, before reaching the base of the cell (Fig. 3*D* and Fig. S1*A*). In some rare cases, pollen tubes even grew upward in the *ktn1-5* mutant and appeared blocked at the tip of the papilla (Fig. S1*B*). To test the direct impact of stigma CMTs on pollen tube growth direction, we examined whether the destabilization of CMTs in Col-0 papillae could affect pollen tube growth. To this end, we treated stigmas by local application of the depolymerizing microtubule drug oryzalin in lanolin pasted around the style. After 4 hours of drug treatment, no more CMT labelling was detected in papillae, while CMTs were clearly visible in mock-treated (DMSO) stigmas (Fig. 4*A*). Stigmas were then pollinated with Col-0 pollen and one hour later, pollen tubes were found turning on drug-treated but not on control papillae (Fig. 4*B* and *C*). However, the number of coils was significantly lower than on *ktn1-5* papillae. Indeed, 25% of the pollen tubes made at least 2 coils in *ktn1-5* papillae whereas this percentage represented only 4% on the oryzalin treated stigmas (Fig. 3*D* and 4*C*). Altogether, these results confirm that stigmatic CMTs contribute to the directional growth of pollen tubes in papilla cells.

**Fig. 3.**
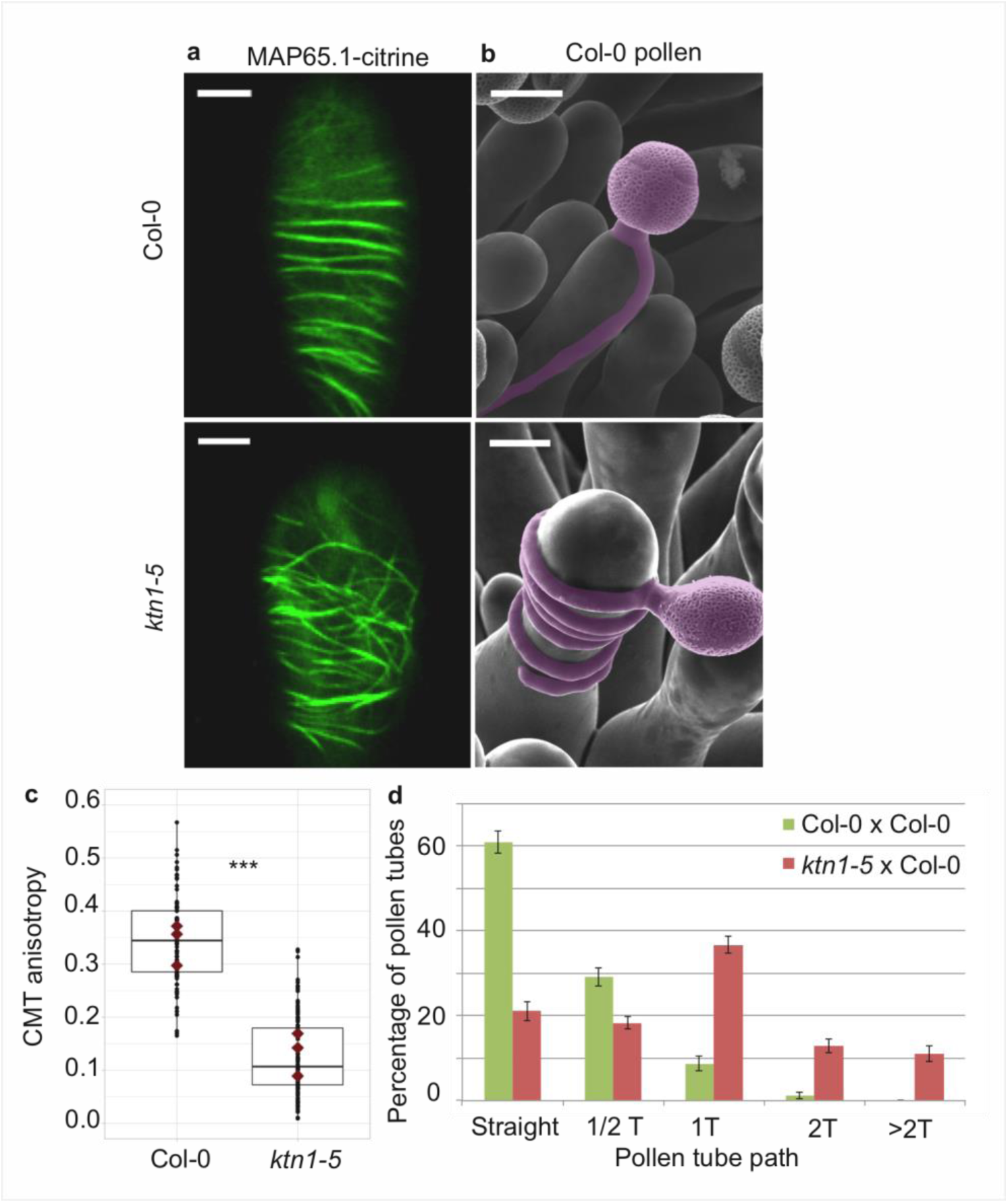
Effect of CMT organisation on pollen tube path. (*A*) Confocal images of papilla cells expressing MAP65.1-citrine in Col-0 and *ktn1-5* at stage 13. Scale bars, 5 μm. (*B*) SEM images of Col-0 and *ktn1-5* papillae pollinated with Col-0 pollen grains; pollen and tubes were artificially colorised. Scale bar, 10 μm. (*C*) CMT anisotropy of Col-0 and *ktn1-5* papilla cells at stage 13. N(Col-0) = 10 stigmas, n(Col-0) = 106 papillae, N(*ktn1-5*) = 11stigmas, n(*ktn1-5*) = 114 papillae. Statistical differences were calculated using a Shapiro-Wilk test to evaluate the normality and then a Wilcoxon test with ***P < 0.01. (*D*) Quantification of the number of turns (T) made by Col-0 pollen tubes on *ktn1-5* and Col-0 papillae. Data are expressed as mean +/-s.e.m. Statistical difference was found between pollen tube path within *ktn1-5* and Col-0 papillae and was calculated using an adjusted Chi-Square test for homogeneity (2 degrees of freedom), ***P < 0.01. N(Col-0) = 27 stigmas, n(Col-0) = 251 papillae, N(*ktn1-5*) = 23 stigmas, n(*ktn1-5*) = 327 papillae.

**Fig. 4.**
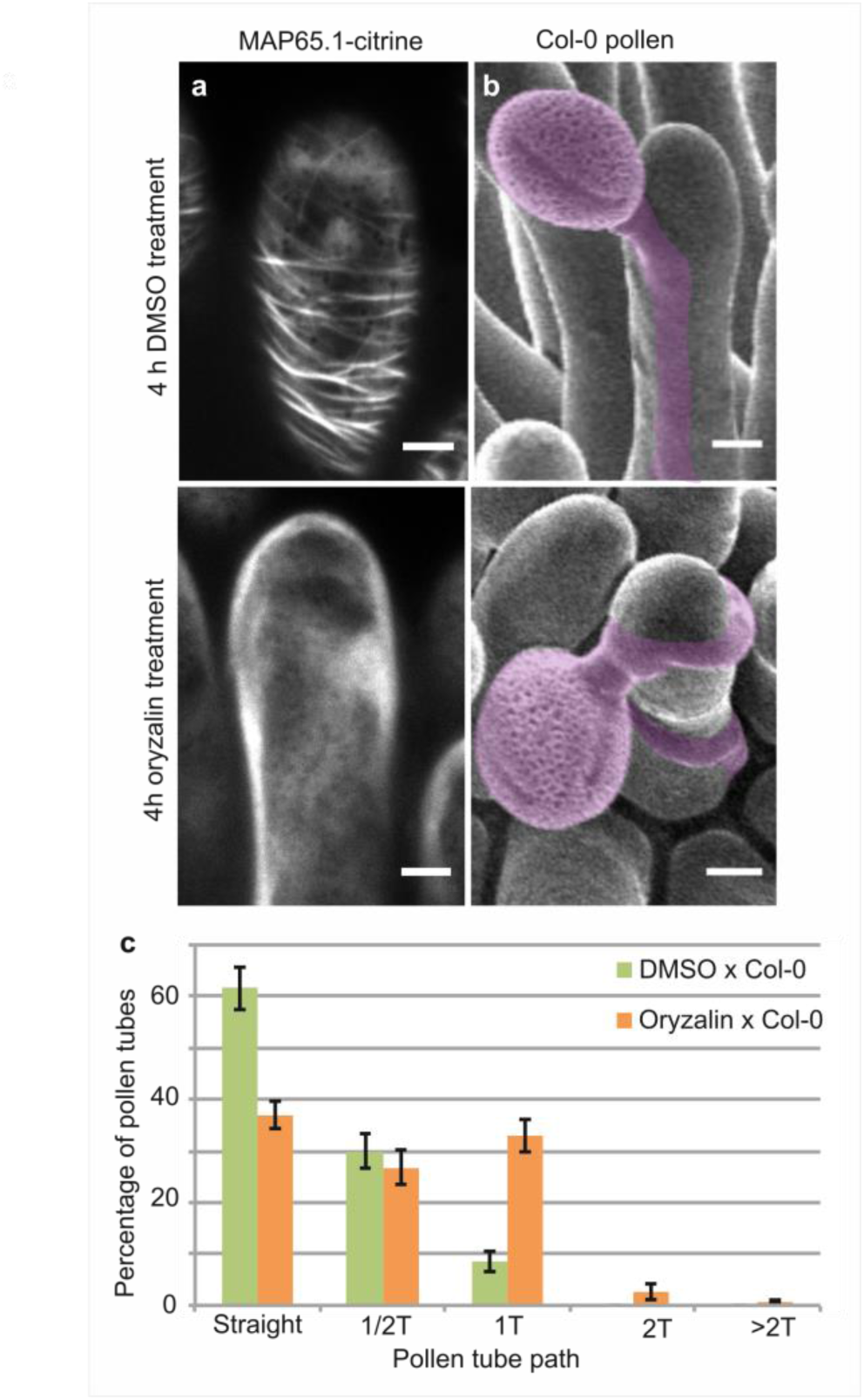
Local oryzalin application on Col-0 stigmas promotes MT destabilization and induces Col-0 pollen tube coils. (*A*) Col-0 papilla cells expressing MAP65.1-citrine after 4 hours of DMSO (top) or oryzalin (bottom) local treatment. (*B*) SEM images of DMSO-(top) or oryzalin-treated (bottom) Col-0 stigmas pollinated with Col-0 pollen grains; pollen and tubes were artificially colorised. (*A* and *B*) Scale bars, 5 μm. (*C*) Quantification of the number of turns (T) made by Col-0 pollen tubes on drug-treated and control papillae. Data are expressed as mean +/-s.e.m. Statistical difference was found between pollen tube path within DMSO (control) and oryzalin-treated papillae and was calculated using an adjusted Chi-Square test for homogeneity (2 degrees of freedom), ***P < 0.01. N(DMSO) = 12 stigmas, n(DMSO) = 117 papillae, N(oryzalin) = 16 stigmas, n(oryzalin) = 149 papillae.

### Mechanical properties of the cell wall are disturbed in *ktn1-5* papilla cells

Because the main role of CMTs in plant cells is to guide the trajectory of CSCs, thereby impacting the mechanical anisotropy of the cell wall, we analysed the cell walls of Col-0 and *ktn1-5* papillae following pollination. First, using Transmission Electron Microscopy (TEM), we found Col-0 pollen tubes to penetrate the cuticle and to grow between the two layers of the papilla cell wall, as previously described (Kandasamy et al., 1994), in both Col-0 and *ktn1-5* papilla cells (Fig. 5*A*). We did not detect any significant difference in the ultrastructure of cell walls (Fig. S2). Interestingly, as the pollen tube progresses through the papilla cell wall, it generates a bump (external deformation) and an invagination (internal deformation) in the cell wall. As such deformations could reflect differences in wall properties, we quantified the external (extD) and internal (intD) deformation following pollination of Col-0 and *ktn1-5* stigmas. To visualize more clearly this deformation, we pollinated stigmas expressing the plasma membrane protein LTI6B fused to GFP (LTI6B-GFP) with pollen whose tube was labelled with the red fluorescent protein RFP driven by the ACT11 promoter (Rotman et al., 2005) (Fig. 5*B*). We found that Col-0 pollen tubes grew with almost equal extD and intD values in Col-0 cell wall. However, the ratio between extD and intD was about 3 when Col-0 pollen tubes grew in *ktn1-5* papilla cells (Fig. 5*B*-*D*). These quantitative data were consistent with the observation that pollen tubes appeared more prominent on *ktn1-5* stigmatic cells using SEM (Fig. 3*B* and Fig. S1). Assuming that stigmatic cells are pressurized by their turgor pressure, this may reflect the presence of softer walls in *ktn1-5* papillae. To test this hypothesis, we assessed the stiffness of Col-0 and *ktn1-5* papilla cell walls using Atomic Force Microscopy (AFM) with a 200 nm indentation. We found that cell wall stiffness in *ktn1-*5 papilla cells was about 30% lower than that in stage 13 WT cells (Fig. 5*E* and *F*). To confirm this result, we investigated the stiffness of the papilla cell wall on WT stigmas at stage 15, where increased coiled pollen tubes were detected (Fig. 2*A* and *B*). We found the cell wall to be softer than that of papillae at stage 13 but stiffer than that of the *ktn1-5* (Fig. 5*E* and *F*). We then reasoned that the presence of softer walls should also affect the pollen tube growth rate, stiffer walls reducing growth. Thus, measuring the growth rate would provide an indication of the resistance force encountered by the tube while growing in the papilla cell wall. We thus monitored the growth rate of Col-0 pollen tubes in Col-0 and *ktn1-5* papillae. We found that pollen tubes grew faster (∼ x 1.8) within *ktn1-5* papillae (Fig. 5*G*). Similarly, we found that pollen tube growth on stage 15 papillae was faster (∼ x 1.6) than on stage 13 papillae (Fig. 5*H*). These results suggest that the coiled phenotype is associated with a higher pollen tube growth rate and hence, that papilla cell walls of *ktn1-5* and stage-15 papillae exhibit less resistance to tube penetration. Altogether, these data suggest that KATANIN-dependent mechanical properties of papilla cell wall play a role in pollen tube guidance.

**Fig. 5.**
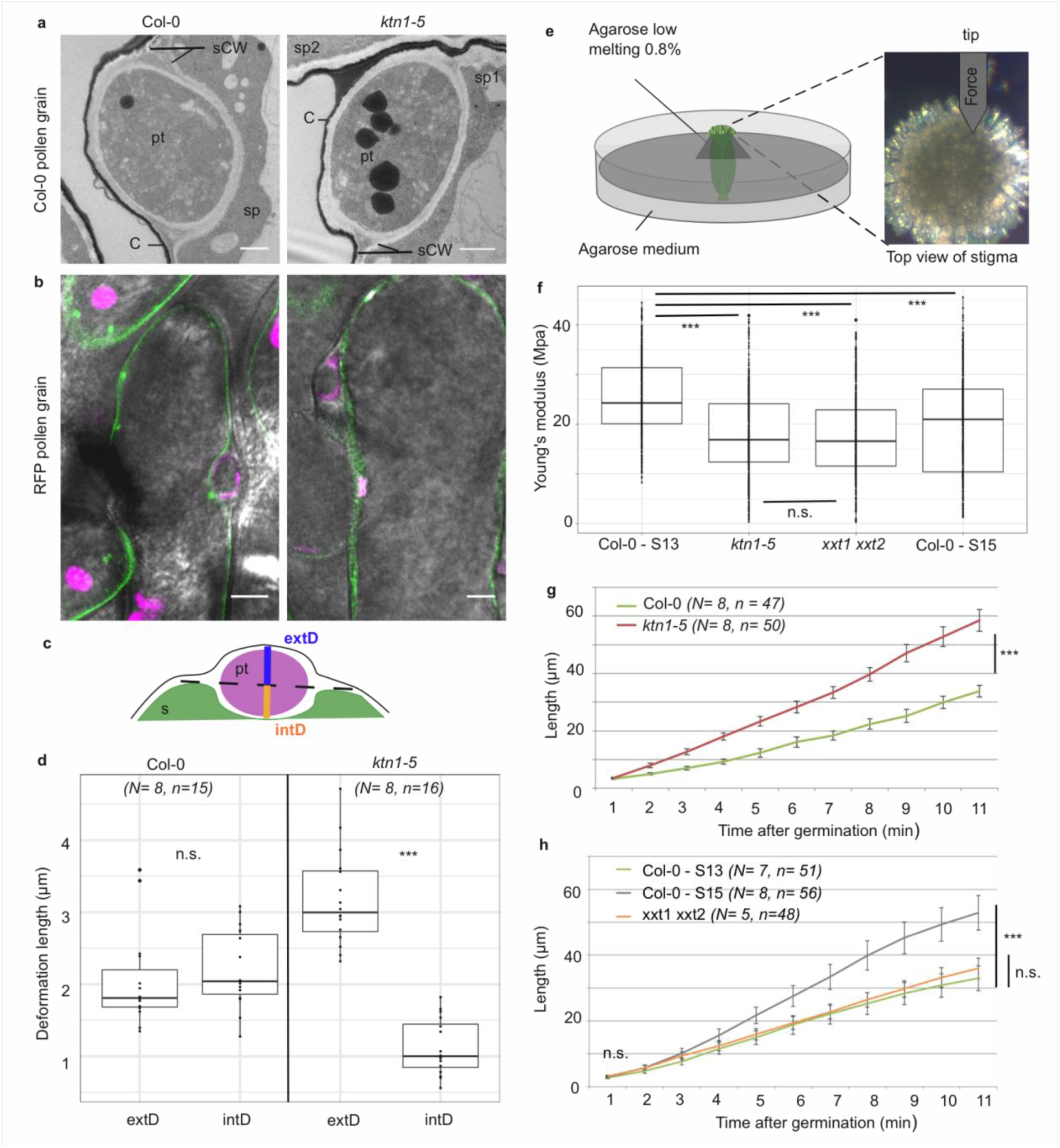
Mechanical properties of papilla cells. (*A*) Location of a Col-0 pollen tube in the cell-wall of Col-0 and *ktn1-5* papillae by TEM. pt = pollen tube, C = stigma cuticle, sCW = stigma cell wall, sp = stigma papilla. Scale bar, 1 μm. (*B*) Confocal images of Col-0 and *ktn1-5* papillae expressing the plasma membrane marker LTI6B-GFP pollinated with an RFP-expressing pollen. Scale bar, 5 μm. (*C*) Diagram showing the procedure used for evaluating the external (extD) and internal (intD) deformations made by Col-0 pollen tubes. (*D*) External and internal deformations caused by Col-0 pollen tube growth in Col-0 and *ktn1-5* papillae. (*E*) Drawing of the AFM experimental setup. Dissected pistils were inserted in agarose medium and fixed with low melting agarose for measurements. (*F*) Young’s modulus values of the papilla cell wall for Col-0 at stage 13 (N = 4 stigmas, n = 8 papillae), *ktn1-5* (N = 5 stigmas, n = 9 papillae), *xxt1 xxt2* (N = 4 stigmas, n = 11 papillae) and Col-0 at stage15 (N = 4 stigmas, n = 10 papillae. (*G*) Mean of travel distances made by Col-0 pollen tubes in Col-0 and *ktn1-5* papillae. (*H*) Mean of travel distances made by Col-0 pollen tubes in papillae of Col-0 at stage 13, Col-0 at stage 15, *xxt1 xxt2* at stage 13. *D-E* and *G-H*, Statistical differences were calculated using a Shapiro-Wilk test to evaluate the normality and then a T-test. *** P < 0.01, n.s. = non significant. For *H*, we found a significant difference (***P < 0.01) between Col-0 at stage 13 and 15, but non significant (n.s.) between Col-0 at stage 13 and *xxt1 xxt2* at stage 13.

### Defects in papilla cell-wall composition are not sufficient to affect Col-0 pollen tube behaviour

The *katanin1/fra2* mutant was initially described as a mutant impaired in cell wall biosynthesis and CMT array organisation (Burk et al., 2001). This prompted us to investigate whether other mutants affected in cell wall biogenesis might exhibit the coiled pollen tube phenotype. Indeed, in the simplest scenario, the presence of softer walls, whatever the cause, may be sufficient to induce extra coiling due to faster pollen tube growth. We selected mutants impaired in the cellulose synthase complex (*kor1.1, prc1* and *any1*), hemicellulose biosynthesis (*xxt1 xxt2, xyl1.4*) and pectin content (*qua2.1*) (Table S1). Strikingly, none of the 6 cell wall mutants displayed the coiled pollen tube phenotype (Fig. S3). This suggests that the relation between CMT organisation, cell wall stiffness and pollen tube trajectory is stricter than anticipated. Because CMTs guide cellulose deposition, they also control the directional elongation of plant cells (Baskin, 2005). Indeed, *katanin1/fra2* cells are wider and shorter than wild type (16, 18), and we may predict that the shape of papilla cells in the mutant might be similarly altered. The contribution of stigmatic CMTs to pollen tube growth may thus be mediated by papilla cell shape only; for instance, wider cells in *ktn1-5* would promote pollen tube coiling. To check that possibility, we measured the length and width of papilla cells in Col-0 (at stage 13 and 15), *ktn1-5, xxt1 xxt2* and *any1* (Fig. S4). We did not find any correlation between papilla length and the coiled phenotype. Indeed, coiled phenotype was observed in stage-15 WT papillae that were longer than those at stage-13, as well as in *ktn1-5* papillae that had a length similar to stage-13 WT papillae. The correlation between papilla width and coiled phenotype was also not clear-cut. As anticipated, *ktn1-5* mutant exhibited wider papillae than Col-0. Stage-15 WT papillae were also significantly larger than stage-13 WT papillae, where coils were observed. However, *xxt1 xxt2* and *any1* papillae were also wider than the WT but did not display the coiled phenotype. Altogether, this suggests that papilla morphology is not sufficient to explain the coiled phenotype. Because CMT disorganisation in papillae affects both wall stiffness and mechanical anisotropy, we next investigated the relative contribution of these two parameters in pollen tube growth. We focused our analysis on the *xxt1 xxt2* double mutant. Indeed, this mutant was reported to display CMT orientation defects and reduced stiffness of the cell wall in hypocotyl cells, when compared with the WT (Xiao et al., 2016). As the pollen tube phenotype on *xxt1 xxt2* stigmas was similar to the WT, we wondered whether the cell wall stiffness of stigmatic cells was actually affected in these mutant papillae. Using AFM, we found the papilla cell wall of the mutant to be about 30% softer than that of Col-0 papillae, i.e. very similar to that of *ktn1-5* (Fig. 5*F*). However, despite this similarity, the pollen tube growth rate in *xxt1 xxt2* stigmas was identical to Col-0 stigmas at stage 13 (Fig. 5*H*). In addition, contrary to *ktn1-5*, pollen tubes were not prominent while growing in *xxt1 xxt2* papilla cell walls (Fig. S3*G*). These results suggested that although *ktn1-5* and *xxt1 xxt2* papillae had similar cell wall stiffness as inferred from AFM measurements, they exhibited different mechanical constraints to pollen tube growth. Because of the close relationship between CMT organisation, CMF deposition and cell wall rigidity (Xiao et al., 2016; Xiao and Anderson, 2016), we suspected that CMT and CMF organisation might be different between the two mutants and be the possible causal agent of stiffness differences and the related tube coiled phenotype observed in *ktn1-5*. To check this possibility, and to gain insight into the predominant cellulose pattern in the stigmatic cell walls, we stained CMFs by using Direct Red 23. We found cellulose fibres to be highly aligned and slanted to the longitudinal axis of the papilla in stage 13 Col-0 stigmas, whereas *ktn1-5* and stage 15 Col-0 papillae both displayed a clearly disordered CMF pattern (Fig. 6a) with CMFs forming thicker and more spaced bundles (Fig. 6b). By contrast, CMFs in *xxt1 xxt2* papillae organized in a dense and well oriented pattern of fibres, mostly perpendicular to the papilla long axis, which resembled Col-0 stage 13 CMF organisation (Fig. 6a and b). These results suggest that KTN1 loss-of-function alters mechanical properties of the papilla cell wall in a complex manner, lowering both its stiffness to pressure applied perpendicularly to its surface (e.g., by the AFM indenter) and its resistance to pollen tube penetration. This latter modification appears as the main cause of the pollen tube coiled phenotype. By contrast, loss of XXT1 and XXT2 functions, by perturbing cell wall stiffness to indenter pressure only, does not induce change in pollen tube growth. Altogether, our data suggest that the mechanical anisotropy of papilla cell walls plays a key role in pollen tube trajectory.

**Fig. 6.**
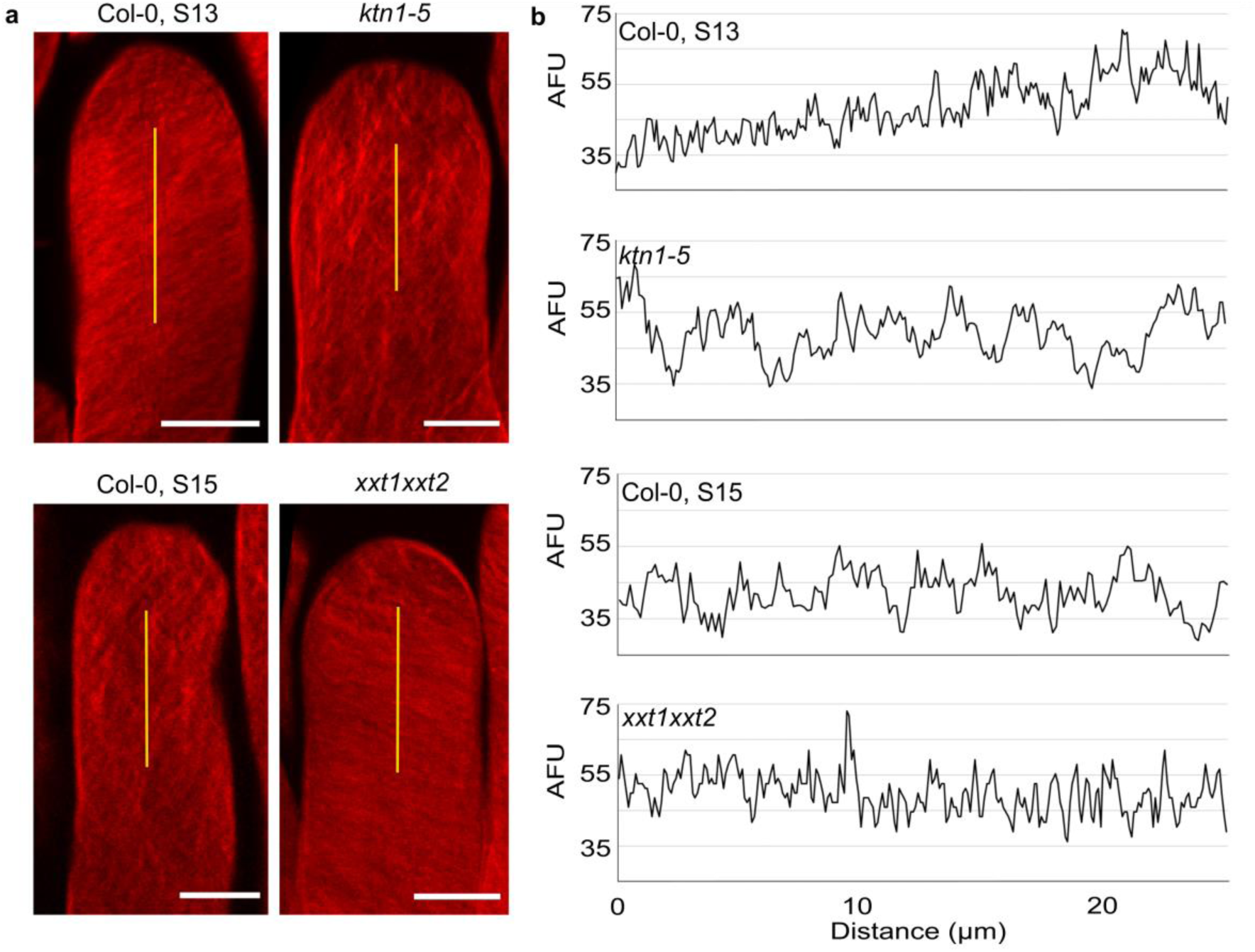
Cellulose microfibril organisation in Col-0 stage 13, *ktn1-5*, Col-0 stage 15 and *xxt1 xxt2* papillae. (*A*) 3D-projections of z-stack images of Col-0 stage 13, *ktn1-5*, Col-0 stage 15 and *xxt1 xxt2* papilla cells stained with Direct Red 23 dye. Scale bars, 10 µm. (*B*) Plot profiles of the fluorescence intensity (AFU, Arbitrary Fluorescence Units) along the yellow lines. Note the similarities of profiles between Col-0 stage 13 and *xxt1 xxt2* on one hand, and between *ktn1-5* and Col-0 on the other hand.

## DISCUSSION

How the pollen germinates a tube and how pollen tube growth is regulated have been the object of many investigations (Mizuta and Higashiyama, 2018; Palanivelu and Tsukamoto, 2012). The use of in vitro pollen germination as well as semi-in vivo fertilization assays, together with the analysis of mutants defective in pollen or ovule functions, have rapidly expanded our knowledge of the mechanisms that sustain pollen tube growth, its guidance toward the ovule and the final delivery of male gametes within the embryo sac (Cameron and Geitmann, 2018; Dresselhaus and Franklin-Tong, 2013; Higashiyama and Yang, 2017). At the cellular level, cytoskeleton has been extensively studied during pollen tube elongation (Fu, 2015), highlighting the critical role played by the actin microfilaments in pollen-tube tip growth through delivery of materials for the biosynthesis of the plasma membrane and cell wall. By contrast, CMTs seem to have a lower importance in this process, drugs affecting CMT polymerisation having no significant effects on pollen-tube growth rate, yet altering the capacity of pollen tubes to change their growth direction (Gossot and Geitmann, 2007). Rearrangements of actin microfilaments (Iwano et al., 2007) and destabilization of CMTs (Samuel et al., 2011) have been described in stigmas following compatible pollination in Brassica species. However, CMT pattern and dynamics during papilla development have never been reported. Our data show that during the course of stigma maturation, which is associated with papilla cell elongation (Fig. S4*A* and *B*), CMT bundles progressively move from perpendicular (anisotropic) to the elongation axis at stage 12 to disorganised (isotropic) at stage 15. This correlates with the known CMT dynamics during elongation in plant cells, where CMT arrays are highly anisotropic in young cells and become more isotropic as cells differentiate (Baskin, 2005; Landrein and Hamant, 2013). The progressive randomisation of CMT orientation we observed in papilla cells is accompanied by an increased coiled growth of pollen tubes in papillae. Similarly, when CMTs are destabilized by the microtubule depolymerizing drug oryzalin, coiled pollen tubes are more frequently observed compared with untreated control stigmas. These results reveal a link between the stigmatic CMT cytoskeleton organisation and the trajectory that pollen tube takes while growing in the papilla cell wall. The fact that the most striking effect on pollen tube growth was found on *ktn1-5* mutant suggests that the coiled phenotype depends not only on the CMT organisation but implicates other factors. Among various phenotypic alterations described in loss-of-function mutants for *KTN1*, are the impaired cell mechanical properties and cell elongation, defects in CMT organisation, cell wall composition and CMF orientation (Burk et al., 2001; Burk and Ye, 2002; Ryden et al., 2003). Interestingly, we found that the protuberance of pollen tubes at the surface of *ktn1-5* papillae was associated with a faster growth rate and a lower rigidity of the cell wall compared with Col-0. This places mechanics of the cell wall as a likely component involved in the coiled phenotype. Surprisingly, of the six cell wall mutants we analysed, including *xxt1 xxt2* and *prc1* known to exhibit both abnormal CMT organisation and softer cell walls, none induced the coiled phenotype displayed by *ktn1-5* papillae. More remarkably, despite the similar stiffness of *xxt1 xxt2* and *ktn1-5* papilla cell walls measured in our AFM experiments, pollen tubes behaved differently on these two stigmas. This indicates that components other than cell wall stiffness are involved in the pollen tube coiled phenotype and that mechanical properties of the cell wall are not identical in the two mutants. Indeed, though sharing a similar alteration of cell wall stiffness, the global plant morphology of these mutants is clearly distinct. The *ktn1-5* mutant was described to have a severe reduction in cell length and an increase in cell width in all organs (Burk et al., 2001). This cellular phenotype was attributed to the distorted deposition of CMFs correlated with the isotropic orientation of CMTs, whereas in WT cells, CMFs like CMTs are oriented perpendicularly to the elongation axis (Burk and Ye, 2002). Similarly, we found that orientation of CMFs is also altered in *ktn1-5* papillae (Fig. 6) and this was associated with larger papilla cells (Fig. S4*C* and *D*). Hence, we may assume that mechanical anisotropy, described as the cell wall anisotropy made by the orientation of the rigid CMFs (Sassi et al., 2014), is impaired in *ktn1-5* papillae. Contrary to the *ktn1-5* mutant, papillae of the *xxt1 xxt2* double mutant display an anisotropic CMF pattern with parallel fibres approximatively oriented transversally to the papilla axis. These data are in agreement with those reported for etiolated hypocotyl cells, where CMFs are largely parallel to one another, straighter than in WT and oriented almost transversely to the long axis of the cell (Xiao et al., 2016). In addition, we found the papilla cell shape of *xxt1 xxt2* stigmas to be similar to Col-0, though with slightly wider papillae (Fig. S3*G* and Fig. S4*E* and *F*). However, *xxt1 xxt2* papillae are less wide than *ktn1-5* papillae, which is consistent with an anisotropic growth of the papillae in the *xxt1 xxt2* mutant compared with *ktn1-5*. Apart from the similar stiffness of papilla cell walls as deduced from AFM measurements, the main differences between *ktn1-5* and *xxt1 xxt2* cell walls are likely to be the cell wall composition, orientation of CMFs and possible changes in molecular connections between cell wall components and plasma membrane and/or cytoskeleton proteins. Indeed, a recent proteomic study revealed that loss of KATANIN function is associated with the decrease in abundance of several cytoskeleton proteins, such as profilin 1, actin-depolymerizing factor 3 and actin 7 (Takáč et al., 2017), whereas targeted quantitative RT-PCR unveiled that several Microtubule-associated protein (MAP) and wall signal receptor genes are downregulated in *xxt1 xxt2* (Xiao et al., 2016). In this latter study, KTN1 expression level was shown to be unchanged compared with Col-0. Altogether, our study suggests that KTN1, by maintaining the papilla mechanical anisotropy, has a key function in mediating early pollen tube guidance on stigma papillae.

The coiled phenotype was not only observed in *ktn1-5* but also in the WT Col-0 papillae at stage 15. Remarkably, several lines of evidence reveal that papilla cells from these two genotypes share common features, such as CMT and CMF increased isotropy, decreased stiffness of the cell wall and less resistance of the wall to pollen tube growth. Isotropic orientation of the cytoskeleton at stage 15 is likely to relate to cell elongation, which is known to be accompanied by cytoskeleton reorganisation (Crowell et al., 2011; Zhang et al., 2014). At the organ level, it has been suggested that the mechanical anisotropy of the wall restrains organ emergence (Sassi et al., 2014). The authors propose that for the same wall stiffness, a cell wall with isotropic properties would lead to larger outgrowth than a wall with anisotropic properties. Our data are consistent with this hypothesis, albeit at the subcellular scale, since large protuberance of papilla wall following pollen tube growth is observed in *ktn1-5* papilla cells, exhibiting walls with isotropic properties.

It remains unclear how mechanical anisotropy guides pollen tube growth. We can hypothesise that as pollen tube grows inside the wall, it encounters recently deposited CMFs on the inner side of the wall (facing the cytoplasm) and older CMFs on the outer side. It is likely that these layers have different mechanical properties related to CMF orientation (Baskin, 2005). Pollen tubes may grow helically by default, as is the case for climbing plants around a cylindrical substrate but the presence of a mechanically reinforced inner wall may slow down and bias the trajectory of the pollen tube. It is worth noting that growth rate is slowed down when pollen tubes pass through a microgap of a microfluiding device, the tubes adapting their invasive force to the mechanical constraints (Nezhad et al., 2013). Based on measurements of pollen tube growth rates, our data show that the tube tip, while progressing in the papilla wall, senses the mechanical features of its environment and reacts accordingly. Hence, it reveals some unanticipated internal and hidden properties of the cell wall. Interestingly, recent work showed that axons were capable of adapting their growth rate and direction according to mechanical constraints, growing straighter on rigid substrate, underlining that mechanical signals are important regulators of pathfinding (Koser et al., 2016).

In the last decades, many studies pointed out that chemical but also mechanical components must be considered to be implicated in pollen tube growth direction (Cameron and Geitmann, 2018). Implication of chemical cues for pollen tube guidance in the stigma remains largely unknown, although the blue copper protein plantacyanin, when overexpressed in Arabidopsis stigmas, was reported to be a possible guidance factor through an as yet undiscovered mechanism (Dong et al., 2005). However, no defect in pollen tube growth directionality was detected in a knock-down plantacyanin mutant, questioning the actual role of this protein as a chemoattractant in Arabidopsis. In our study, we add mechanics as a key player in early pollen tube guidance in the papilla cell. Our results suggest that this role is mediated by a specific CMT/CMF organisation and mechanical anisotropy of the papilla cell, which both are dependent on KTN1. Importantly, these specific mechanical properties of the stigmatic cells prevent emerging pollen tubes to grow upward on papillae and straighten pollen tube direction, helping the tube to find its correct path to the stylar transmitting tract. We uncovered a yet unexpected role in pollen tube guidance for KTN1, which was until now mostly known to be involved in plant development and stress-response regulation. In addition, our study also clearly unveils that the mechanical properties of one single cell (e.g., the stigmatic papilla) impact the behaviour of its neighbouring cell (e.g., the pollen tube).

## MATERIALS AND METHODS

### Plant Materials and Growth Conditions

*Arabidopsis thaliana*, ecotype Columbia (Col-0), *Arabidopsis* transgenic plants generated in this study and *Arabidopsis* mutants were grown in soil under long-day conditions (16 hours of light / 8 hours of dark, 21°C / 19°C) with a relative humidity around 60%. *ktn1-5* (SAIL_343_D12), *xxt1xxt2, prc1.1, qua2.1, xyl1.4, kor1.1* and *any1* mutant lines were described previously (Cavalier et al., 2008; Fagard et al., 2000; Fujita et al., 2013; Lin et al., 2013; Mouille et al., 2007; Nicol et al., 1998; Sampedro et al., 2010; Shoji et al., 2004). All mutants were in Col-0 background except *kor1.1* which was in WS.

### Plasmid construction

We used the Gateway® technology (Life Technologies, USA) and two sets of Gateway®-compatible binary T-DNA destination vectors (Hellens et al., 2000; Karimi et al., 2002) for expression of transgenes in *A. thaliana*. The DNA fragment containing the *Brassica oleracea SLR1* promoter was inserted into the pDONP4-P1R vector. The 165 bp-*LTI6B* fragment was introduced into the pDONR207 vector. *MAP65* gene spanning the coding region from start to stop codons was introduced into the pDONR221 vector. CDS from citrine or GFP were cloned into the pDONP2R-P3 vector. Final constructs, pSLR1::MAP65-citrine and pSLR1::LTI6B-GFP were obtained by a three-fragment recombination system (Life Technologies) using the pK7m34GW and the pB7m34GW destination vectors, respectively. We generated a pAct11::RFP construct by amplifying the promoter of the *A. thaliana Actin 11* gene and cloning it into the pGreenII gateway vector in front of the RFP coding sequence.

### Generation of transgenic lines and crossing

Transgenic lines were generated by *Agrobacterium tumefaciens*-mediated transformation of *A. thaliana* Col-0 as described (Logemann et al., 2006). Unique insertion lines, homozygous for the transgene were selected. We introduced the pSLR1::LTI6B-GFP or pSLR1::MAP65-citrine construct in *ktn1-5* background by crossing and further selecting the progeny on antibiotic containing medium.

### Microscopy

#### Confocal microscopy

Flowers at stages 12 to 15 (Smyth et al., 1990) collected from fluorescent lines were emasculated and stigmas were observed under a Zeiss LSM800 microscope (AxioObserver Z1) using a 40x Plan-Apochromat objective (numerical aperture 1.3, oil immersion). Citrine was excited at 515 nm and fluorescence detected between 530 and 560nm. GFP was excited at 488 nm and fluorescent detected between 500 and 550 nm. RFP was excited at 561 nm and fluorescent detected between 600 and 650 nm.

#### Live imaging

Flowers from stages 12 to 15 were emasculated and pollinated on plants with mature pollen from the pACT11::RFP line. Immediately after pollination, stigmas were mounted between two coverslips. To maintain a constant humidity without adding liquid directly on the stigma surface, we use a wet piece of tissue in contact with the base of the stigma. Pollinated stigmas were observed under a Zeiss microscope (AxioObserver Z1) equipped with a spinning disk module (CSU-W1-T3, Yokogawa) using a 40x Plan-Apochromat objective (numerical aperture 1.1, water immersion). Serial confocal images were acquired in the entire volume of the stigma every 1 µm and every minute. Images were processed with Image J software and pollen tube lengths were measured.

#### Atomic Force Microscopy

Pistils at stage 13 were placed straight in a 2% agar MS medium and 0.8% low-melting agarose was added up to the base of papilla cells. AFM indentation experiments were carried out with a Catalyst Bioscope (Bruker Nano Surface, Santa Barbara, CA, USA) that was mounted on an optical microscope (MacroFluo, Leica, Germany) equipped with a x10 objective. All quantitative measurements were performed using standard pyramidal tips (RFESP-190 (Bruker)). The tip radius given by the manufacturer was 8-12 nm. The spring constant of the cantilever was measured using the thermal tune method and was 35 N/m. The deflection sensitivity of the cantilever was calibrated against a sapphire wafer. All experiments were made in ambient air at room temperature. Matrix of 10×10 measurements (step 500 nm) was obtained for each papilla, with a 1µN force. The Young’s Modulus was estimated using the Nanoscope Analysis (Bruker) software, using the Sneddon model with a < 200nm indentation.

#### Environmental Scanning Electron Microscopy (SEM)

Flowers from stages 12 to 15 were emasculated and pollinated on plants with mature WT pollen. One hour after pollination, pistils were cut in the middle of the ovary, deposited on a SEM platform and observed under Hirox SEM SH-3000 at −20°C, with an accelerating voltage of 15kV. Images were processed with ImageJ software and pollen tube direction was quantified by counting the number of turns made by the tube, only on papillae that received one unique pollen grain.

#### Transmission Electron Microscopy

Stage 13 flowers were emasculated and pollinated on plants with mature WT pollen. One hour after pollination, pistils were immersed in fixative solution containing 2.5% glutaraldehyde and 2.5% paraformaldehyde in 0.1 M phosphate buffer (pH 7.2) and after 4 rounds of 30 min vacuum, they were incubated in fixative for 12 hours at room temperature. Pistils were then washed in phosphate buffer and further fixed in 1% osmium tetroxide in 0.1 M phosphate buffer (pH 7.2) for 1.5 hours at room temperature. After rinsing in phosphate buffer and distilled water, samples were dehydrated through an ethanol series, impregnated in increasing concentrations of SPURR resin over a period of 3 days before being polymerized at 70°C for 18 h, sectioned (65 nm sections) and imaged at 80 kV using an FEI TEM tecnaiSpirit with 4 k x 4 k eagle CCD.

#### Anisotropy estimation

Flowers from MAP65 lines were emasculated and stigmas were observed under confocal microscope. Images were processed with ImageJ software and quantitative analyses of the average orientation and anisotropy of CMTs were performed using FibrilTool, an ImageJ plug-in (Boudaoud et al., 2014). Anisotropy values range from 0 to 1; 0 indicates pure isotropy, and 1 pure anisotropy.

#### Cellulose staining

A droplet of Adigor™ was applied at 2.5% (v/v in water) on the stigma during 4 hours to facilitate subsequent chemical treatments. As described for hypocotyls (Landrein et al., 2013), pistils were incubated in 12.5% glacial acetic acid for 1 hour and dehydrated in 100% and then 50% ethanol for 20 min each. Pistils were then washed in water during 20 min and stored in 1M KOH during 2 days. Pistils were then stained using 0.02% (w/v) Direct Red 23 dye (Sigma-Aldrich) during 4 hours and washed with distilled water. Pistils were observed with a mRFP filter (excited at 561 nm) under a Zeiss LSM 880 confocal microscope using a 63x Plan-Apochromat objective (numerical aperture 1.4, oil immersion). Stigmas were imaged by taking a z-stack of 0.2 μm sections in the papilla cells. Image analysis was performed by doing a 3D-projection and measuring the fluorescence intensity longitudinally along the papilla axis using ImageJ software.

#### Membrane deformation estimation

Flowers from LTI6B lines were emasculated and pollinated with mature pollen from the PACT11::RFP line. 20 minutes after pollination, stigmas were observed under confocal microscope. Serial confocal images every 1 µm encompassing the entire volume of the stigma were recorded and processed with ImageJ software. Plasma membrane deformation was estimated by choosing the slide from the stack that corresponded to the focus plan of the contact site with the RFP-labelled pollen tube. On the Bright field image corresponding to the selected slide, a line was drawn connecting the two ridges of the invagination of the papilla. Two perpendicular lines, one toward the exterior (ExtD) of the papilla to the maximum point of deformation, and the other toward the interior (IntD) on the GFP image were measured, respectively.

### Oryzalin treatment

To avoid contact of pollen grains with liquid, we performed local applications of oryzalin (Chemical service, Supelco) in lanolin pasted (Sassi et al., 2014), around the style, just under the stigmatic cells, at 833 µg/mL (DMSO), for 4 hours at 21°C. Oryzalin-treated pistils were pollinated with mature WT pollen and 1 hour after pollination observed under SEM.

### Statistical analysis

Graph and statistics were obtained with R software or Excel. Statistical tests performed are specified in figure legends.

## ACKNOWLEDGEMENTS

We thank O. Hamant for critical reading of the manuscript and fruitful discussion, A. Boudaoud, S. Bovio and V. Battu for advice on AFM experiments, and the Sice and MechanoDevo team members for discussion, P. Bolland, A. Lacroix and J. Berger for plant care and the PLATIM imaging facility of the SFR Biosciences Gerland-Lyon Sud. We thank the Bordeaux Imaging Centre especially L. Brocard and B. Batailler for TEM microscopy. We also thank the BioMeca® society and Pascale Milani for the AFM measurements. L.R. was funded by a fellowship from the French Ministry of Higher Education and Research. The work was supported by Grant ANR-14-CE11-0021.

## AUTHOR CONTRIBUTIONS

L.R. performed all the experiments and the image analysis, except TEM and AFM, which were subcontracted. F.R. and C.K. contributed to cell imaging setup. L.R., T.G., and I.F.L., designed the study and analysed the data. L.R. and T.G. wrote the manuscript.

## Competing interests

The authors declare no competing interests.

## SUPPLEMENTAL INFORMATION

**Fig. S1.**
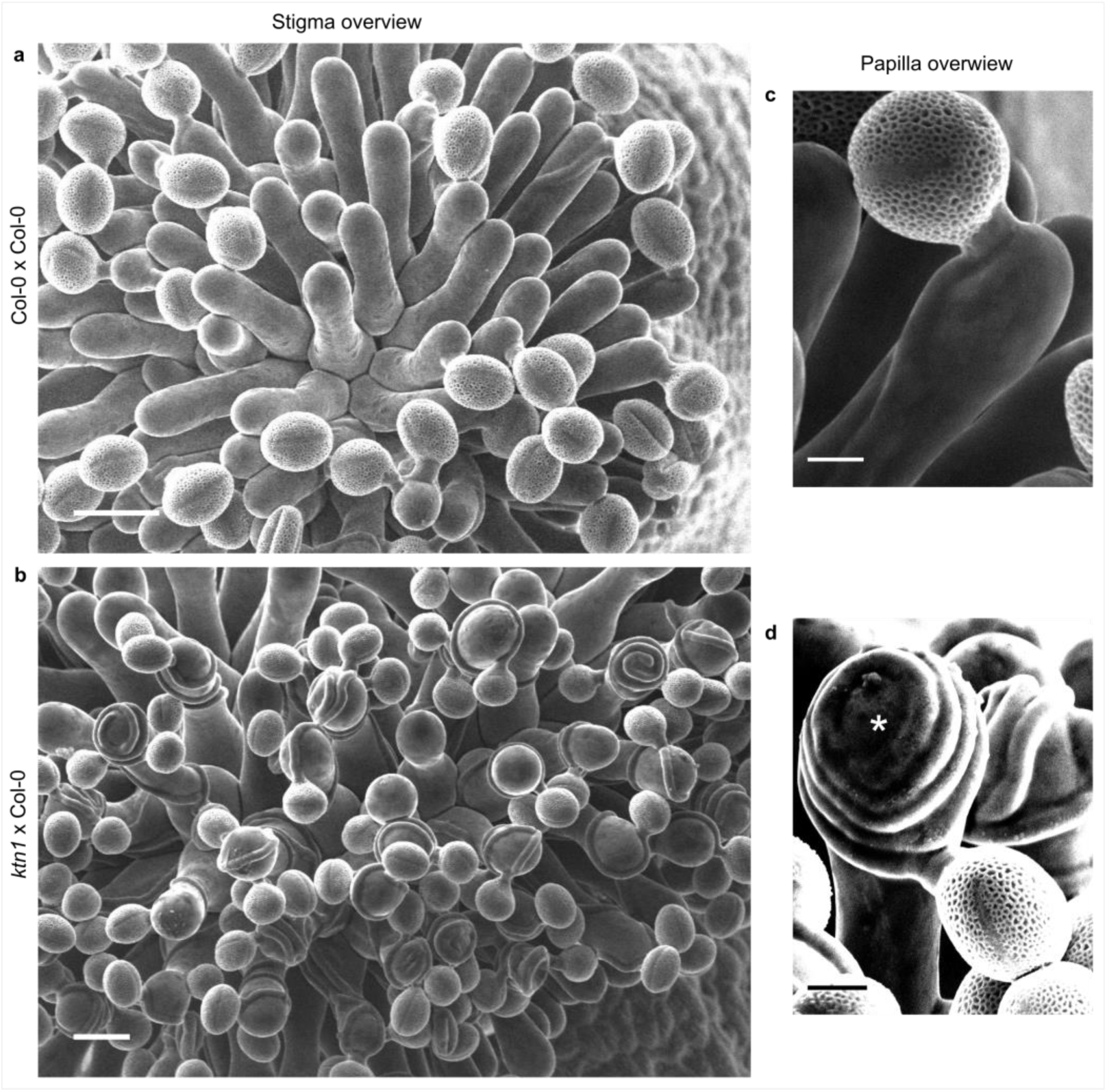
Col-0 pollen tube behaviour on Col-0 and *ktn1-5* papillae at stage 13. (*A* and *B*) Top views of Col-0 (*A*) and *ktn1-5* (*B*) stigmas pollinated with Col-0 pollen grains. Scale bars, 20 µm. (*C* and *D*) Magnification of pollinated papilla cells. (*C*) Col-0 pollen tube grows mainly straight to the direction of the ovules on Col-0 papillae whereas (*D*) it coils around and can even grow upward on *ktn1-5* papilla cells (*). Scale bars, 5 µm.

**Fig. S2.**
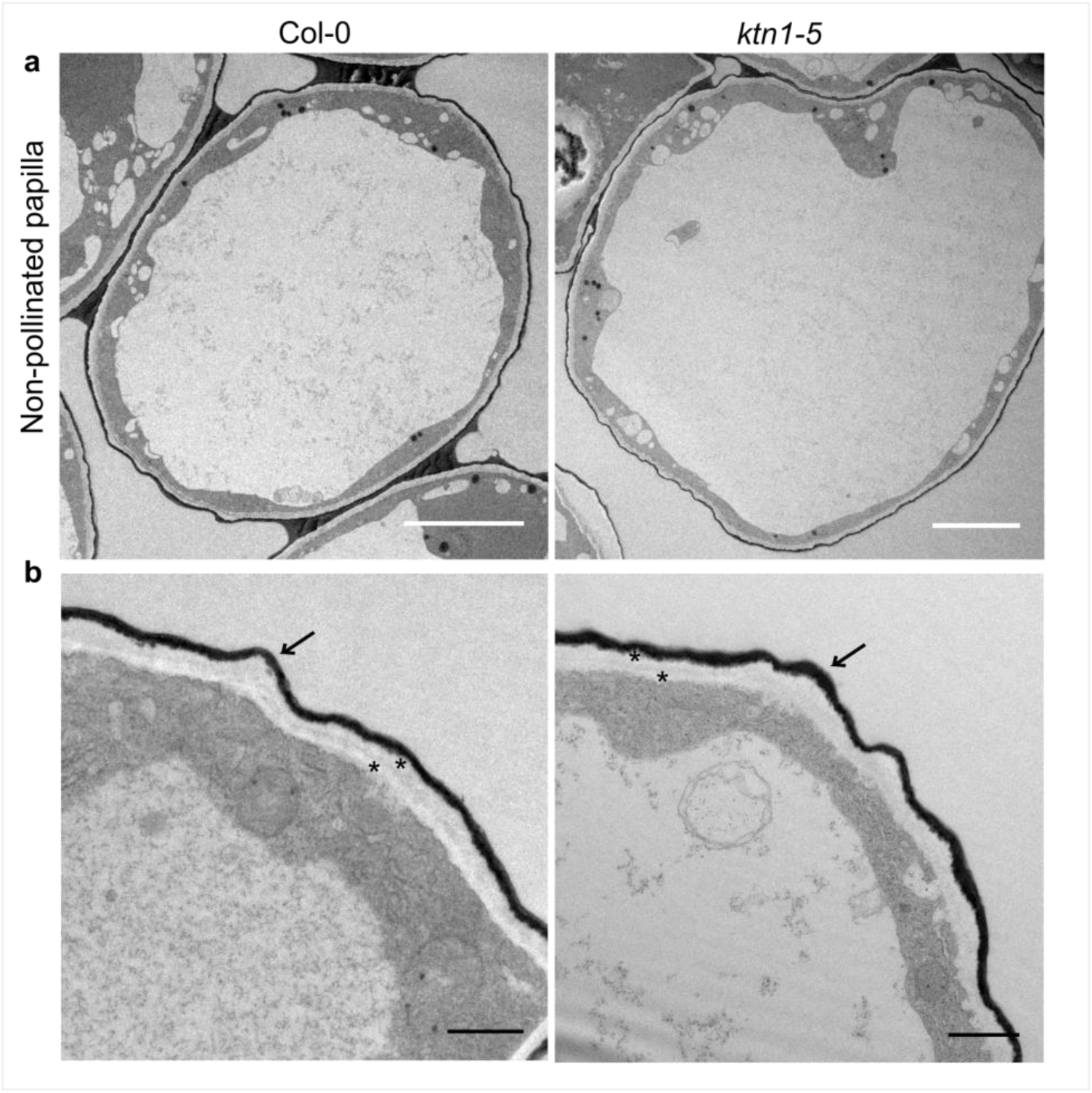
Ultrastructure of Col-0 and *ktn1-5* papilla cells. (*A*) TEM images of non-pollinated Col-0 and *ktn1-5* papillae. Scale bar, 5 μm. (*B*) Both Col-0 and *ktn1-5* papillae show a similar two-layered cell wall (*) delimited by a more electron dense thin layer (arrows) corresponding to the cuticle. Scale bar, 1 µm.

**Fig. S3.**
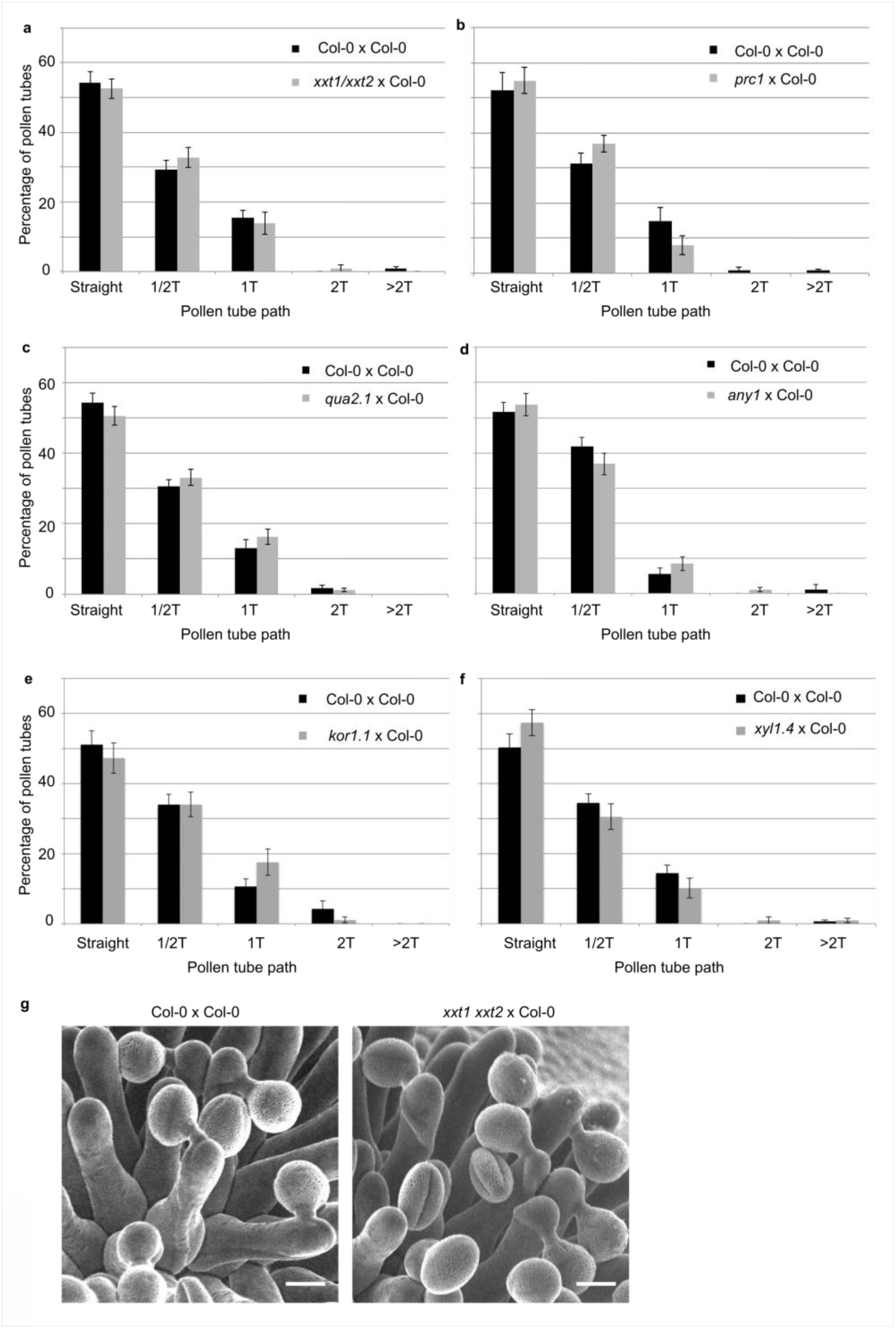
Quantification of the number of coils made by Col-0 pollen tubes on papillae from cell-wall mutants at stage 13. (*A*) *xxt1 xxt2* x Col-0. N(Col-0) = 14 stigmas, n(Col-0) = 116 papillae, N(*xxt1 xxt2*) = 12 stigmas, n(*xxt1 xxt2*) = 116 papillae. (*B*) *prc1* x Col-0. N(Col-0) = 14 stigmas, n(Col-0) = 115 papillae, N(*prc1*) = 12 stigmas, n(*prc1*) = 100 papillae. (*C*) *qua2.1* x Col-0. N(Col-0) = 22 stigmas, n(Col-0) = 160 papillae, N(*qua2.1*) = 18 stigmas, n(*qua2.1*) = 162 papillae. (*D*) *any1* x Col-0. N(Col-0) = 14 stigmas, n(Col-0) = 91 papillae, N(*any1*) = 14 stigmas, n(*any1*) = 95 papillae. (*E*) *kor1.1* x Col-0. N(Col-0) = 14 stigmas, n(Col-0) = 141 papillae, N(*kor1.1*) = 11 stigmas, n(*kor1.1*) = 91 papillae. (*F*) *xyl1.4* x Col-0. N(Col-0) = 18 stigmas, n(Col-0) = 136 papillae, N(*xyl1.41*) = 16 stigmas, n(*xyl1.4*) = 108 papillae. Data are expressed as mean +/-s.e.m. No statistical difference was found between pollen tube path within the cell wall mutants and Col-0 papillae based on an adjusted Chi-Square test for homogeneity (2 degrees of freedom). (*G*) SEM images of Col-0 and *xxt1 xxt2* stigmas pollinated with Col-0 pollen grains. Scale bar, 10 µm.

**Fig. S4.**
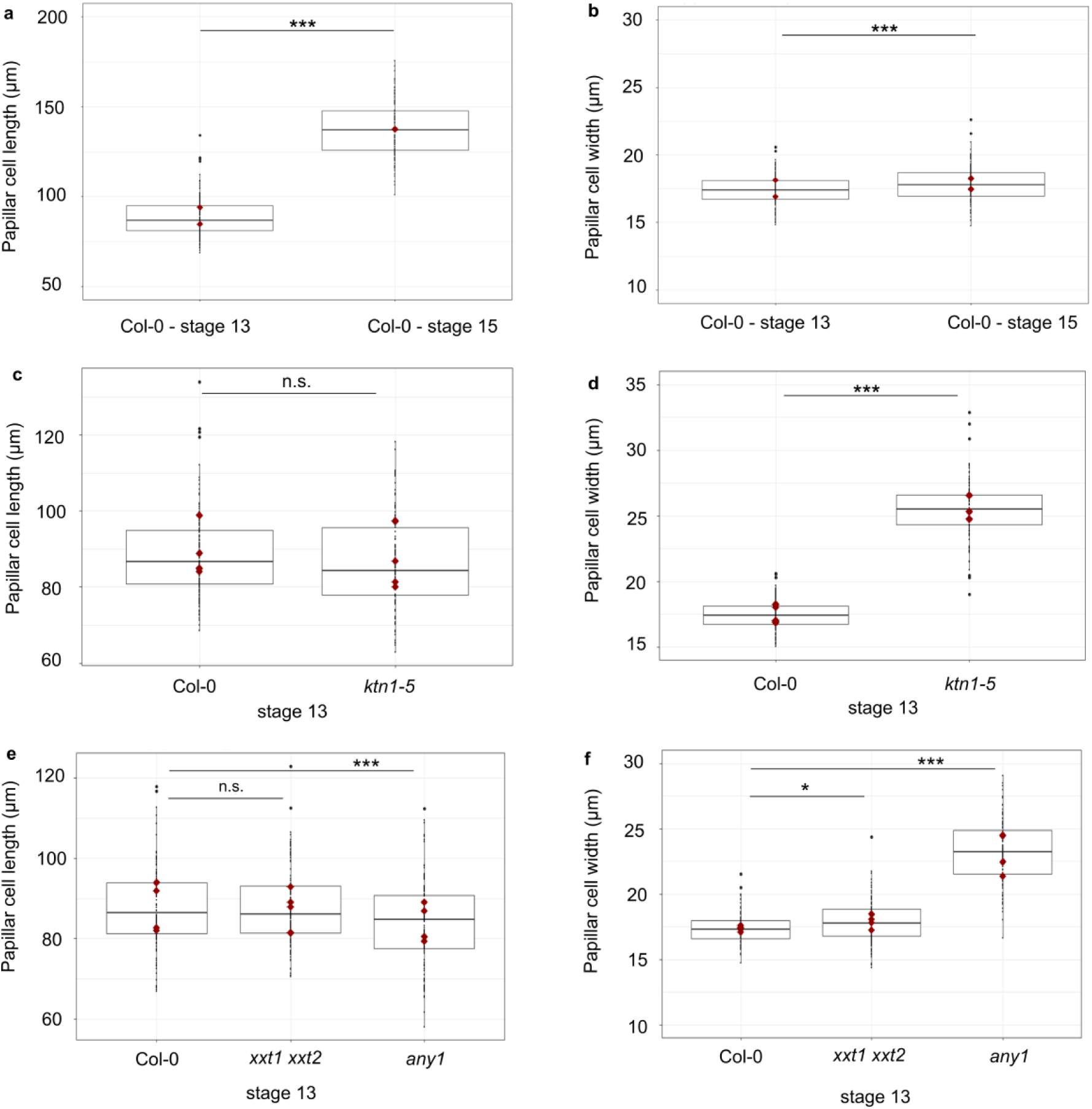
Size of the papilla cells of Col-0 and cell wall mutants. (*A*) Papilla cell length of Col-0 at stage 13 and stage 15. (*B*) Papilla cell width of Col-0 at stage 13 and stage 15. (*C*) Papilla cell length of Col-0 and *ktn1-5* at stage 13. (*D*) Papilla cell width of Col-0 and *ktn1-5* at stage 13. (*E*) Papilla cell length of Col-0, *xxt1 xxt2* and *any1* at stage 13. (*F*) Papilla cell width of Col-0, *xxt1 xxt2* and *any1* at stage 13. Statistical differences were calculated using a T-test. * P < 0.05, *** P < 0.01, n.s. = non significant. N=4 stigmas and n=120 papillae for each genotypes.

**Table S1.**
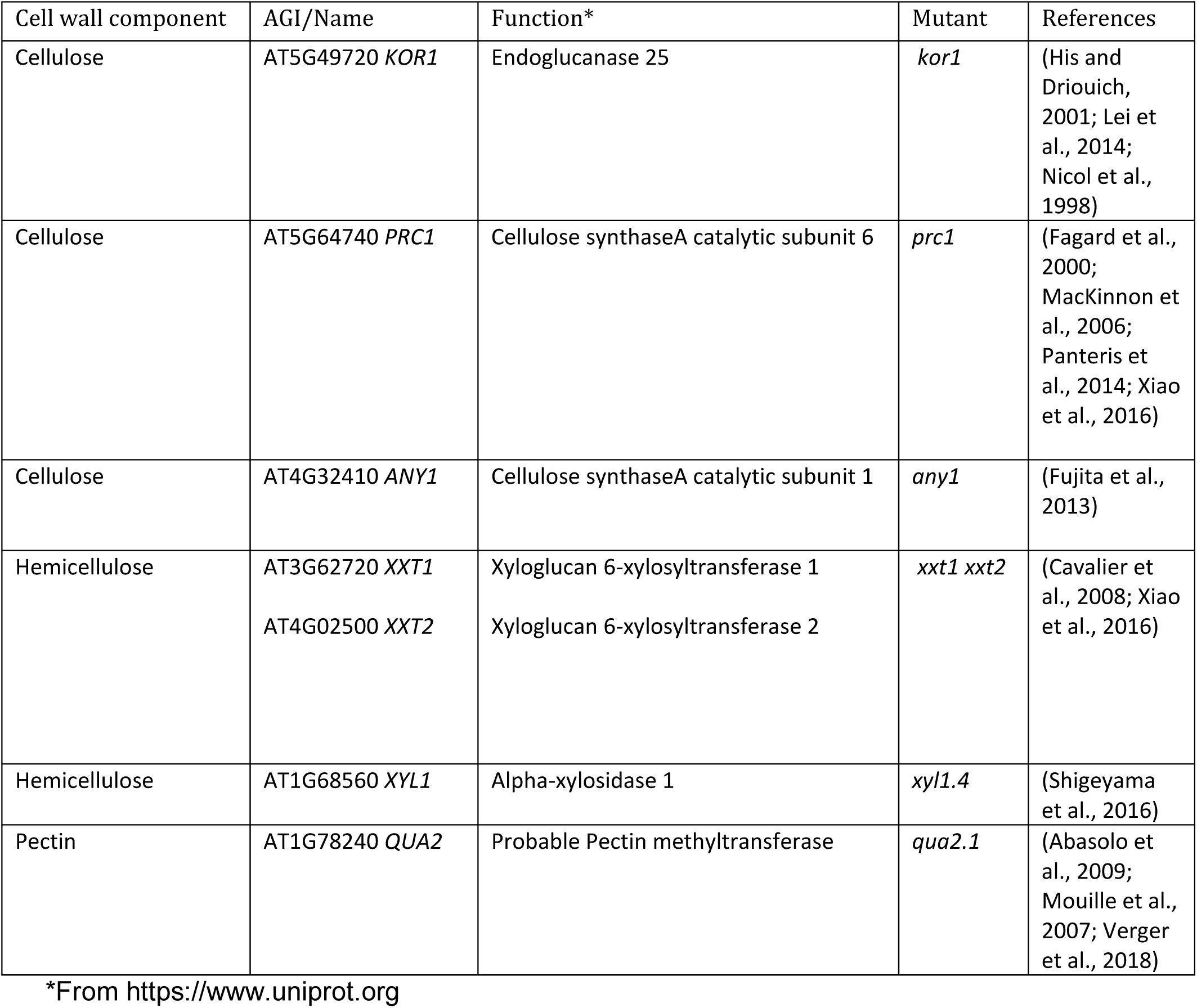
Cell-wall mutants tested.

